# Atomic model of microtubule-bound tau

**DOI:** 10.1101/267153

**Authors:** Elizabeth H. Kellogg, Nisreen M.A. Hejab, Simon Poepsel, Kenneth H. Downing, Frank DiMaio, Eva Nogales

## Abstract

Tau is a developmentally regulated protein found in axons, whose physiological role is to stabilize and bundle microtubules (MTs). Hyper-phosphorylation of tau is thought to cause its detachment from MTs and subsequent aggregation into pathological fibrils that have been implicated in Alzheimer’s disease pathogenesis. Despite its known MT binding role, there is no consensus regarding which tau residues are crucial for tau-MT interactions, where on the MT tau binds, and how binding results in MT stabilization. We have used cryo-EM to visualize the interaction of different tau constructs with MTs at high resolution (3.2-4.8 Å) and used computational approaches to generate atomic models of tau-tubulin interactions. Our work shows that the highly conserved tubulin-binding repeats within tau adopt very similar structures in their interactions with the MT. Each tau repeat binds the MT exterior and adopts an extended structure along the crest of the protofilament (PF), interacting with both α- and β-tubulin, thus stabilizing the interface between tubulin dimers. Our structures agree with and explain previous biochemical data concerning the effect of phosphorylation on MT affinity and lead to a model in which tau repeats bind in tandem along a PF, tethering together tubulin dimers and stabilizing longitudinal polymerization interfaces. These structural findings could establish a basis of future treatments aiming at the selective stabilization of tau-MT interactions.

## Main Text

The microtubule (MT) cytoskeleton is essential for many eukaryotic cellular functions, including cellular trafficking and chromosome segregation. MTs are formed by the head-to-tail assembly of αβ-tubulin dimers into protofilaments (PFs), which in turn associate laterally to form hollow tubes. The properties and cellular functions of MTs are regulated by their interaction with a myriad of MT associated proteins (MAPs). In neurons, MTs interact with a family of “classical” MAPs, including MAP-2, MAP-4, and tau, that contribute to MT organization and stability and are critical to neuronal growth and function. Tau, which constitutes more than 80% of neuronal MAPs, localizes to axons where it functions to stabilize and bundle MTs(*1*) and is developmentally regulated(*2*).

Tau and its homologs are intrinsically disordered proteins containing an array of conserved MT-binding sequences. Full-length tau from adult human neurons(*3*) includes a projection domain, a central, MT-binding region containing four imperfect sequence repeats, R1-4, and a C-terminal domain (Fig. 1A). Different MT binding repeats have been shown to bind to and stabilize MTs(*4, 5*), with the affinity and MT stabilization activity increasing with the number of repeats(*5, 6*). Neurodegenerative disorders known as tauopathies, including Alzheimer’s disease, develop when tau is abnormally phosphorylated and its affinity for MTs is reduced(*7–9*). Dissociation from MTs ultimately leads to the formation of intracellular, filamentous tau aggregates, called neurofibrillary tangles.

**Fig. 1.**
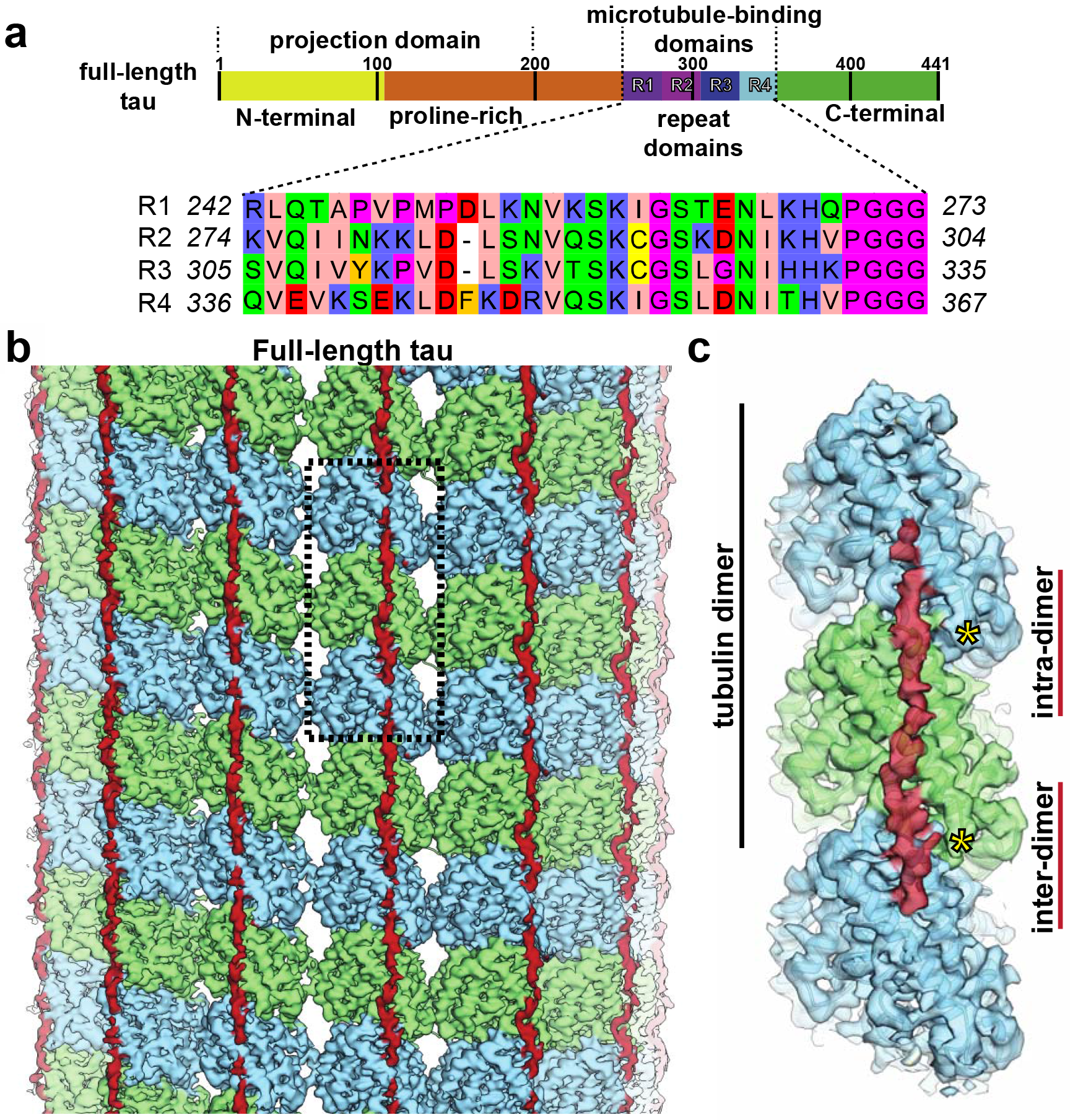
Tau binding to microtubules. **(A)**Schematic of tau domain architecture and assigned functions. The MT-binding domain of four repeats includes residues 242-367. Inset shows the sequence alignment of the four repeat sequences, R1-4, that make up the repeat domain **(B)** Cryo-EM density map (4.1 Å overall resolution) of a MT decorated with full-length tau. Tau (red) appears as a nearly continuous stretch of density along PFs (α-tubulin in green, β-tubulin in blue). **(C)** The footprint of a continuous stretch of tau spans over three tubulin monomers, binding both across intra- and inter-dimer tubulin interfaces (only one repeat of tau is shown for clarity). Position of the C-termini of tubulin indicated with yellow asterisks.

While the high-resolution structure of the amyloid tau aggregates has recently been described(*10*), the physiologically relevant conformation of MT-bound tau is still poorly understood. NMR and fluorescence spectroscopy data indicated that the MT-binding domain of tau could adopt a helical structure upon association with MTs(*11, 12*), while solid-state NMR studies suggested a structured hairpin conformation(*13*). Cryo-electron microscopy (cryo-EM) studies also resulted in contradictory proposals concerning the tau binding site on MTs. While tau was reported to bind along PFs when added to pre-formed, Taxol-stabilized MTs(*14*), it was proposed to bind near the Taxol-binding site on the luminal MT surface if tubulin was co-polymerized with tau in the absence of the drug(*15*). The low resolution attainable at the time of these reports added to the uncertainty of their conclusions. Defining the high-resolution structure of MT-bound tau has implications for Alzheimer’s treatment through a better understanding of how tau phosphorylation and mutations contribute to loss of native function and onset of disease. Here, we present atomic structures of MT-bound tau using a hybrid approach that combines state-of-the-art cryo-EM imaging with computational protein structure determination.

In order to visualize tau on MTs, we first assembled dynamic MTs in the presence of excess full-length tau, and carried out cryo-EM studies following our previously established protocols to obtain high-resolution MT structures(*16–18*). The cryo-EM reconstruction (overall resolution of 4.1 Å (Fig. S1C)) shows tau as a narrow, discontinuous density along each PF (Fig. 1B), approximately following the outermost ridge on the MT surface defined by the H11-H12 helices of both α-and β-tubulin. The tau density is adjacent to the site of attachment of the unstructured C-terminal tails (Fig. 1C), in agreement with biochemical data highlighting the importance of the acidic tails of tubulin for tau affinity(*19, 20*). This location is fully consistent with the previous cryo-EM reconstruction by Al-Bassam and Milligan(*14*), but the higher resolution now allows us to conclude that the density correspond to a fully extended chain, rather than an α-helical segment, as previously proposed. To probe for the existence of a tau-MT binding site within the MT interior, we added tau either to pre-formed MTs or to polymerizing tubulin, both in the absence of Taxol (see Methods). None of the reconstructions showed any tau density on the MT luminal surface (data not shown). Thus, contrary to a previous report(*15*), we identified a single binding site on the MT exterior regardless of preparation conditions.

In addition to full-length tau, we also examined two N- and C-terminally truncated tau constructs that included either all four repeats (4Repeat) or just the first two repeat sequences (2Repeat) (Fig. S1A and B, respectively). The 4Repeat- and 2Repeat-MT cryo-EM reconstructions are indistinguishable from the full-length tau at the present resolution (estimated overall resolutions of 4.8 Å and 5.6 Å, respectively (Fig. S1C)). In all three cases the length of the density ascribed to tau corresponds to an extended chain of about 27 aa (see later), with a connecting region of very weak density that would accommodate another 3-4 extended residues, thus adding up to the length of the tau repeats (31 aa). This finding strongly suggests that each segment of the observed density is attributable to a single repeat. The fact that the 2Repeat, 4Repeat, and full-length tau reconstructions are basically indistinguishable indicates that the repeats likely adopt similar structures on the MT-surface upon binding. It is important to note that, given the nature of our reconstruction procedures (see Methods), the cryo-EM structures for these tau constructs correspond to an average of the different repeats.

The resolution for the tau density within these reconstructions (4.6-6.5 Å) is significantly lower than that for tubulin (4 to 4.5 Å) (Fig. S2), which could be due to sub-stoichiometric binding, flexibility, compositional heterogeneity (i.e. sequence differences between the repeats), or a combination of these factors. Thus, we decided to pursue the structure of a synthetic tau construct comprising four identical copies of R1 (R1×4) (Fig. 2). The cryo-EM reconstruction of the R1×4-MT reached an overall resolution of 3.2 Å (Fig. 2A, Fig. S1C), with local resolution for tau ranging from 3.7 Å to 4.2 Å (Fig. S2). Again, tau appears as regularly spaced segments separated by fragments of more discontinuous density. Consistent with conclusions from previous studies(*5, 21*), the alternation of well-defined and weaker tau density indicates that tau has tightly-bound segments interspersed with more mobile regions. We built a poly-alanine model into the best-resolved segment of the R1×4 tau density, at the inter-dimer interface, that accommodated 12 residues. Prior studies showed that tau fragments as small as 18 amino acids were sufficient to promote MT polymerization and stabilization(*4*). The tau binding site partly overlaps with that of the motor domain of kinesin (Fig. S3), consistent with reports that tau binding interferes with kinesin-MT attachment(*22*).

**Fig. 2.**
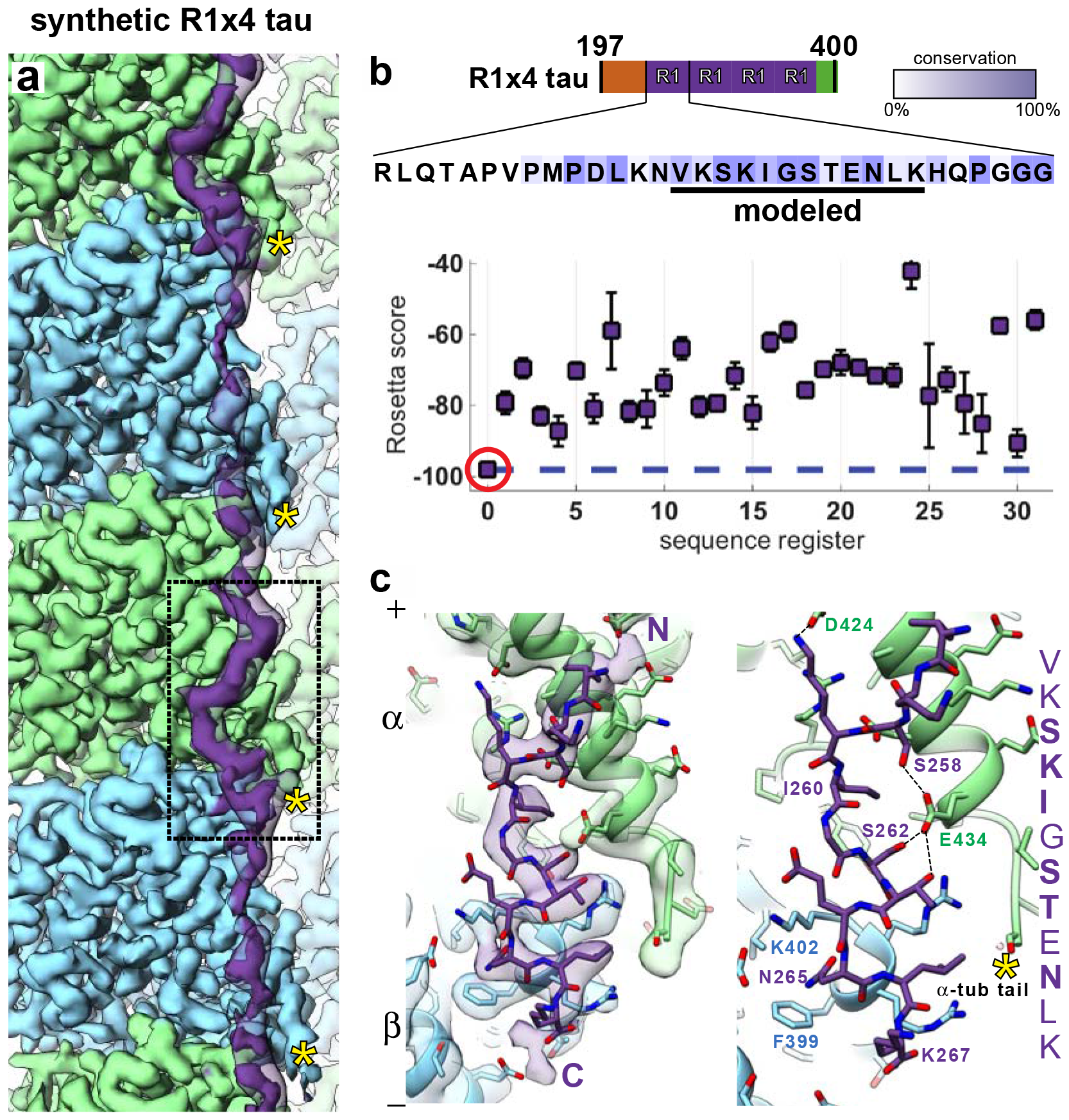
Near-atomic resolution reconstruction of synthetic R1×4 tau on microtubules. **(A)**In the 3.2 Å cryo-EM reconstruction of tau-bound microtubules, the best-ordered segment of tau is bound at the interface between tubulin dimers (boxed) (lower threshold density for tau is shown in transparency). **(B)** Rosetta modeling reveals a single energetically preferred sequence register (red circle) for the best-ordered tau region, corresponding to a conserved (underlined) 12-residue stretch of residues within the R1 repeat sequence. Intensity of colored boxes indicates extent of conservation among tau homologs. **(C)** Atomic model of tau and tubulin, with (left) and without (right) the density map, showing the interactions over the inter-dimer region. C-termini of tubulin indicated with yellow asterisks in a and c.

Due to the lack of unambiguously identifiable large sidechains and the absence of regular secondary structure, obtaining the correct sequence register for tau proved challenging. Therefore, we used Rosetta(*23*) as an unbiased approach to simultaneously assess the energetics and density-agreement of potential tau sequence registers (see Methods). We identified a single register and conformation that was significantly favored, both energetically and based on fit to density, and that included aa 256-267 (VKSKIGSTENLK) (Fig. 2B). An alternative refinement protocol(*24*) allowing further conformational sampling converged to the same amino acid sequence and structure (Fig. S4). The modeled segment corresponds to a conserved 12 aa sequence motif that lies within the 18 aa fragment previously shown to be sufficient to promote MT polymerization *in vitro*(*4, 25*). In the proposed atomic model, conserved residues play critical roles in MT-tau interactions (Fig. 2C). Ser258 and Ser262 form hydrogen bonds with α-tubulin Glu434. Importantly, phosphorylation of the universally conserved Ser262 (Fig. S5) has been previously shown to strongly attenuate tau-MT binding(*26*) and is a marker of Alzheimer’s disease(*27*). Though Thr263 is also positioned to hydrogen-bond with Glu434, hydrophobic substitutions are tolerated at this position (Fig. S5), indicating that this interaction may not be essential. Conserved Lys259 is positioned to interact with an acidic patch on α-tubulin formed by Asp424, Glu420, and Glu423. Ile260, conserved in hydrophobic character across all R1 sequences, is buried within a hydrophobic pocket formed by α-tubulin residues Ile265, Val435, and Tyr262 at the inter-dimer interface (Fig. 2C). Asn265, universally conserved amongst repeats in classical MAPs (Fig. S5), forms a stabilizing intramolecular hydrogen bond within the type II’ β-turn formed by residues 263-266. Finally, Lys267 is positioned to interact with the acidic α-tubulin C-terminal tail, and its basic character is conserved.

It was not possible to reliably model the peptide backbone beyond this 12-residue stretch of the R1 tau density. We hypothesized that residues 242-255 lack ordered interactions with tubulin. This region is rich in prolines in R1, in contrast to the distinct, conserved hydrophobic pattern present in both R2 and R3 (Fig. 1A, S5). To further investigate whether the repeats can form additional interactions with the MT surface, we next obtained a cryo-EM reconstruction of a synthetic tau construct containing four copies of repeat R2 (R2×4) (overall resolution of 3.9 Å) (Fig. 3A and S1C). Although the densities for R1×4 and R2×4 are similar at the inter-dimer interface (Fig. 2 and S6A), the R2×4 reconstruction has additional tau density along the surface of β-tubulin. We modeled a stretch of 27 backbone residues into the R2×4 tau density, spanning three tubulin monomers and corresponding almost to a length of approximately 80 Å, the length of a tubulin dimer repeat on the MT lattice. Using the same Rosetta-guided modeling as with R1, we refined all possible tau sequence registers. As with R1, we discovered a single, strongly preferred register and conformation for R2 (Fig. 3B) that was similarly robust when modeling with a more computationally intensive sampling protocol. Importantly, the solutions for R1 and R2 corresponded to the same sequence register. At equivalent positions at the inter-dimer interface, the R1 and R2 atomic models are virtually identical (Fig. 3C), with conserved residues sharing critical tau-tubulin contacts. For two non-conserved positions, Cys291(R2) instead of Ile260(R1), and Lys294(R2) instead of Thr263(R1) (Fig. 3C), the nature of the interactions are preserved (free cysteines demonstrate strong hydrophobic character(*28*) and Lys294 likely interacts with the acidic C-terminal tail (Fig. 4)). The robustness of this solution, with the same sequence register and atomic details from two independent maps, provides very high confidence that our sequence assignments – and the resulting atomic details – are correct. The consensus register of our atomic models for R1 and R2 is further supported by previous gold-tagging results, which localized MAP2 Lys368 (Fig. S6) to the H12 helix in tubulin at the interface between tubulin subunits(*14*). In our atomic model, the equivalent Gln276 of R2 maps to the intra-dimer interface at the C-terminal end of helix H12 in β-tubulin (Fig. S6), in excellent agreement with the labeling studies.

**Fig. 3.**
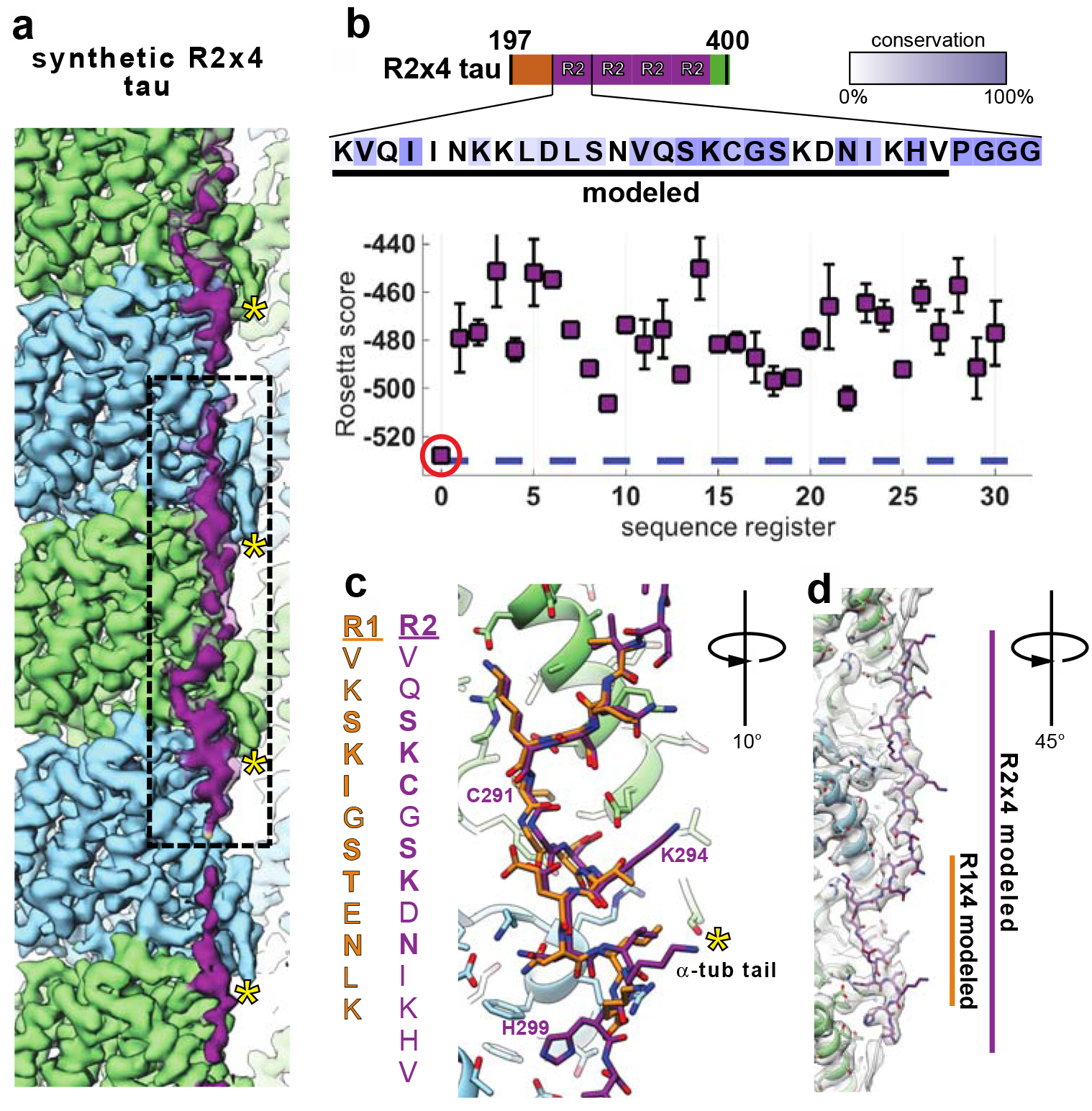
High-resolution reconstruction of synthetic R2×4 tau on microtubules. **(A)** The (3.9 Å) cryo-EM reconstruction of R2×4 is highly similar to that of R1×4 but reveals a longer stretch of ordered density for tau along the MT surface (lower threshold density for tau is shown in transparency). **(B)** Rosetta modeling supports a sequence register (red circle) for the R2 sequence binding to tubulin analogous to that for R1 and shown in Fig. 2. **(C)** Major tau-tubulin interactions at the inter-dimer cleft are highly similar between the R1 (shown in goldenrod for easier visualization) and R2 (purple) sequences. **(D)** Extending the model to account for the additional density reveals an almost entire repeat of tau, spanning three tubulin monomers (centered on α-tubulin and contacting β-tubulin on either side) with an overall length of ~80 Å (the approximate length of a tubulin dimer). C-termini of tubulin indicated with yellow asterisks in a and c.

**Fig. 4.**
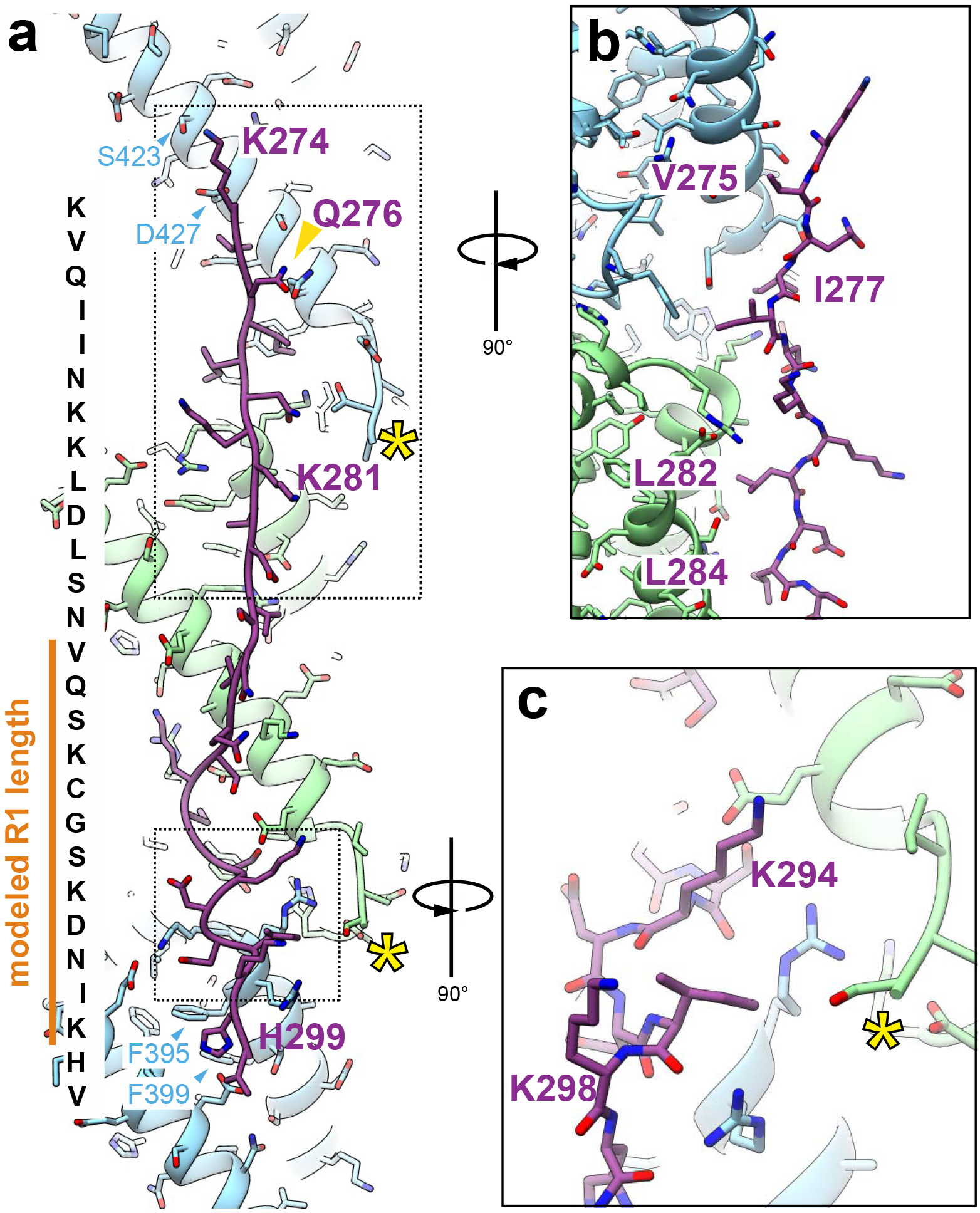
Repeat 2 makes additional conserved interactions with the MT surface. **(A)** Atomic model of the R2 repeat, shown along with the corresponding R2 sequence. The position of previously studied gold-labeled tau(*14*) is indicated with a gold arrowhead (see also Fig. S6). Boxed out regions show: **(B)** the hydrophobic packing of tau residues on the MT surface, and **(C)** the positioning of two R2 lysines with potential interactions with the α-tubulin acidic C-terminal tail.

Additional residues in our R2×4 model include the peptide VQIINKK (not conserved in R1), which had been previously shown to promote MT polymerization(*25*) and to bind independently of the conserved sequence that targets the inter-dimer interface(*6*). This R2 peptide localizes to the intra-dimer interface and is sufficiently close to interact with the β-tubulin C-terminal tail (Fig. 4B). Residues Val275, Ile277, Leu282 and Leu284 (Fig. 3D, 4B) are buried against the MT-surface and tolerate only conservative hydrophobic substitutions in R2, R3 and R4 (Fig. S5). As further support for our model, Lys274 and Lys281 of tau have previously been shown to be crucial for tau-MT binding(*6*), and in our atomic model Lys274 is sufficiently close to interact with an acidic patch on the MT-surface formed by Asp427 and Ser423 (Fig. 4A), while Lys281 is well positioned to interact with the β-tubulin C-terminal tail (whereas Lys280, which had little effect on tau affinity, is not) (Fig. 4A). Finally, the highly conserved His299 in R2 is buried in a cleft formed by β-tubulin residues Phe395 and Phe399 (Fig. 3C and Fig. 4A).

We observe that the major tau binding site between tubulin dimers localizes to the previously defined ‘anchor point’, a tubulin region that remains unaltered during the structural changes accompanying nucleotide hydrolysis within the MT, and even when comparing assembled and disassembled states of tubulin(*16*). Indeed, superimposing the bent (4I4T(*29*)) and straight forms of tubulin (3JAR(*16*)) (Fig. 5, right) reveals that the tau-tubulin contacts at the tubulin dimer interface are significantly preserved in the former. This finding agrees with and explains the observation that tau added to tubulin under non-assembling conditions results in the formation of tubulin rings(*30*), and with the co-purification of classical MAPs, including tau, through temperature-driven tubulin assembly-disassembly cycles.

**Fig. 5.**
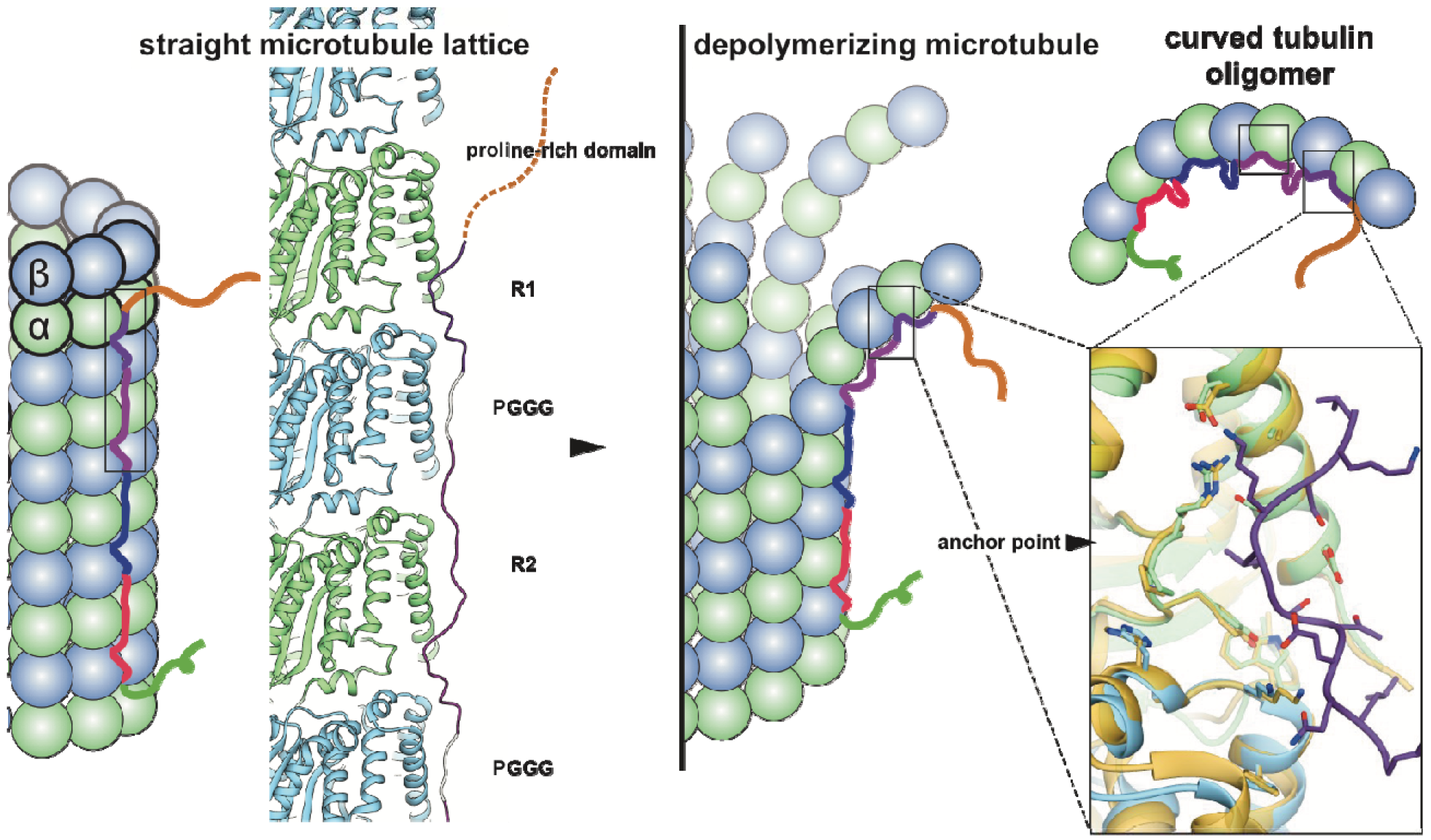
Model of full-length tau binding to microtubules and tubulin oligomers. Our structural data leads to a model of wild-type tau in which the four repeats bind in tandem along a MT protofilament. Notice that we do not observe strong density for the region that would correspond to the PGGG motif, which is modeled in gray for illustrative purposes and must be highly flexible. Tau binding at the inter-dimer interface, interacting with both α-and β-tubulin, promotes association between tubulin dimers. The tau binding site is also the location of the previously identified “anchor point” (right-most box, gold is bent tubulin, PDB:4I4T (*29*) and blue/green is straight tubulin, PDB: 3JAR (*16*)), and thus tau-tubulin interactions are unlikely to change significantly with protofilament peeling during disassembly (center) or when bound to small, curved tubulin oligomers (top-right).

Although we could model most of the residues in a tau repeat, we could not clearly visualize the highly conserved PGGG motif, which would correspond to the region connecting the modeled segments. A prior study reported a β-turn structure for the PGGG motif(*13*). However, the construct utilized in that study contained PGGG at the N- and C-termini, and it is possible that the reported state is present only when there is no binding of PGGG-flanking residues. Recent findings suggest that the PGGG motif may play a role when bound to unassembled tubulin(*31*).

Tau is an important cellular factor required for specialized MT arrangements, and it has been long debated how tau may cross-link MTs to form higher order structures. Our results suggest that all four tau repeats are likely to associate with the MT surface in tandem through adjacent tubulin subunits along a PF. This modular structure explains how alternatively spliced variants (regulated developmentally or in disease states) can have essentially identical interactions with tubulin but different affinity based on the number of repeats present. However, if one or two of the repeats were used as “spacers”, then tau would possibly be able to reach across adjacent PFs, as previously proposed(*32, 33*), or even cross-link MTs, using its repeat sequences. Additionally, other sequence elements distinct from the repeats, such as the projection domain, could engage with tubulin to add tethering points with the MT surface, both within and between MTs. Such additional interactions of tau would likely engage the unstructured tubulin C-terminal tails, as otherwise we would likely have seen some difference between our full-length and shorter tau construct reconstructions.

In summary, our MT-tau structures lead to a model in which each tau repeat spans both intra-and inter-dimer interfaces, centered on α-tubulin and connecting three tubulin monomers. In the context of full-length tau, repeats are likely to bind one after the other in tandem along a PF, promoting MT polymerization and stabilization by tethering tubulin dimers together across longitudinal interfaces (Fig. 5, left). Other structural elements could potentially be involved in inter-PF or inter-MT interactions. Regardless of the construct used, we observe an extended conformation for the MT-binding repeat. The details of the repeat structure and MT interactions are supported by our extensive modeling on two independently determined reconstructions, and account for previous biochemical and structural observations. However, a number of important questions remain open concerning the range of configurations of full-length tau-MT binding under physiological, non-saturating conditions. The strikingly conserved PGGG motif is not visible in any of our reconstructions and its structural role remains unsolved. It is possible that the flexibility of this motif is needed to allow binding to the different conformational states of tubulin in oligomers and MTs. Alternatively, the PGGG motif may play a critical role in association of tau with substrates other than tubulin, such as actin(*34*).

## Acknowledgements

We thank Abhiram Chintangal and Paul Tobias for computational support. We also thank Patricia Grob and Daniel Toso for EM support. We thank Basil Greber for help with initial model building.

## Author contributions

Conceptualization,E.H.K., N.M.H., and E.N.; Methodology, E.H.K., N.M.H., F.D., and E.N.; Investigation, E.H.K., N.M.H., F.D., and E.N.; Writing-Original Draft, E.H.K. and E.N.; Writing-Review and Editing, all authors; Funding acquisition, E.N.; Resources, S.P.; Supervision, E.N.

## Data and materials availability

Atomic models are available through the PDB with accessions codes XXXX, all cryo-EM reconstructions are available through the EMDB with accession codes XXXX. All data and code is available upon request.

